# Spatiotemporal secondary hair follicle development in the Lanyu pig (Sus scrofa taivanus): a novel pelage hair follicle model

**DOI:** 10.64898/2026.04.26.720264

**Authors:** Chein-Hong Lin, Ting-Yung Kuo, Yuan-Yu Hsueh, Shy-Jou Sheih, Ming-Jer Tang, Chia-Ching Wu, Lynn L.H. Huang, Michael W Hughes

## Abstract

Large full-thickness (LFT) skin wounds remain a major clinical challenge, and progress in regenerative medicine has been limited by poor translation from animal models to humans. A key limitation is that commonly used species such as mice, rats, and rabbits are loose-skinned, whereas humans are tight-skinned with distinct skin architecture. Although pigs more closely resemble human skin, widely used breeds have lost secondary (vellus-like) hair follicles through artificial selection, restricting their utility for studying ectodermal organ regeneration.

Here, we characterize the development, patterning, and molecular features of secondary hair follicles in the Lanyu pig (*Sus scrofa taivanus*), an indigenous breed that retains these structures. Whole-mount and histological analyses revealed two distinct follicle populations: primary follicles arranged in stable triplet clusters and smaller secondary follicles distributed interstitially. A developmental time course using alkaline phosphatase (ALP) staining identified sequential stages of secondary follicle morphogenesis—placode, hair germ, hair peg, and mature follicle—occurring after primary follicle establishment.

Immunohistochemical analysis demonstrated conserved epithelial– mesenchymal interactions, progressive epithelial stratification, and dynamic β-catenin signaling during secondary follicle development. Keratin expression patterns and follicular architecture closely resembled those of human vellus hair follicles, supporting the translational relevance of this model. Notably, secondary follicles were retained into adulthood, and genetic analyses of outcrossed animals suggest that this trait follows an autosomal dominant inheritance pattern.

Together, these findings establish the Lanyu pig as a tight-skinned mammalian model that preserves vellus-like hair follicles, providing a platform for investigating hair follicle–mediated skin regeneration and improving translational relevance for human wound healing.

## INTRODUCTION

Large full-thickness (LFT) skin wounds represent a major clinical challenge and remains a significant source of morbidity. Despite extensive advances in regenerative medicine, most findings derived from animal models fail to translate effectively to human patients, largely due to fundamental differences in skin architecture and physiology^1^. In particular, commonly used laboratory species including mice, rats, and rabbits are loose-skinned, whereas human skin is tight-skinned, resulting in distinct wound healing dynamics and regenerative capacities^2^.

Hair follicles are essential components of skin architecture and play critical roles in tissue homeostasis and repair. During pelage development, hair follicles are broadly classified into primary and secondary types^3, 4^. Primary follicles develop first, are larger, and exhibit complex structures including dermal papillae, sebaceous glands, and arrector pili muscles. In contrast, secondary follicles develop later, are smaller, and typically lack associated arrector pili muscles^5, 6^. Importantly, secondary follicles are structurally and functionally analogous to human vellus hair follicles, which predominate in human torso skin.

Hair follicles contribute to wound healing through multiple mechanisms, including sebaceous secretion, structural support, and activation of hair follicle stem cells (HFSCs)^7^. Upon injury, HFSCs migrate into the wound bed and differentiate into multiple epidermal lineages^8, 9^. However, in humans, LFT wound healing results in scar formation without regeneration of ectodermal appendages such as hair follicles and glands, leading to permanent loss of HFSCs and function resulting in reduced tissue resilience.

Porcine skin closely resembles human skin in both structure and immune composition, positioning it as a valuable model for translational research. Compared to rodents, pigs exhibit similar epidermal thickness, dermal architecture, and immune cell populations, including keratinocyte-associated Toll-like receptors, Langerhans cells, dendritic cells, and macrophages^10, 11^. However, more commonly utilized porcine models including Landrace and Red Duroc pigs have undergone artificial selection that suppresses secondary hair follicle development, limiting their utility for studying vellus hair biology and regeneration^3, 12-14^.

This limitation is critical, as human skin predominantly contains vellus hair follicles, which do not regenerate following LFT injury. Notably, although loose-skinned animal models including the classical laboratory mouse regenerate only secondary follicles after wounding, and rare species such as the African spiny mouse can regenerate both primary and secondary follicles, no tight-skinned mammalian model has been shown to regenerate these structures to our knowledge^15-17^.

Therefore, an ideal translational model should combine tight skin architecture with the presence of secondary hair follicles in order to study skin regeneration after LFT wound healing. The Lanyu pig (*Sus scrofa taivanus*), an indigenous Taiwanese breed, represents a potential solution. In this study, we characterize the development, patterning, and molecular features of secondary hair follicles in the Lanyu pig, establishing a model system that more closely recapitulates human skin architecture and provides a platform for studying vellus hair follicle biology and regenerative wound healing.

## MATERIAL & METHODS

### Fetal and adult Lanyu minipig model

All experiments were approved by the Institutional Animal Care and Use Committee (IACUC) of National Cheng Kung University, College of Medicine. Lanyu pigs were purchased from the Taitung Animal Propagation Station, Livestock Research Institute, Taitung Taiwan. The animals were maintained in a climate-controlled environment with normal light/dark cycles. Adult pregnant female Lanyu pigs were anesthetized with isoflurane under veterinarian supervision. The dorsal skin of embryonic, stages for fetal age day 89 (e89), and fetal age day 100 (e100), were harvested. After euthanasia with sodium pentobarbital, the dorsal skin of a juvenile neonate Lanyu piglet day 1 (p1), a 6-month-old adult Lanyu pig, a 6-month-old adult Lee-Sung pig, and a 6-month-old adult Mitsai pig were harvested at the Livestock Research Institute, Tainan Taiwan. All animals were included in the study unless deemed unhealthy by the onsite veterinarian.

### Human skin samples

The human adult skin samples were obtained with the appropriate approvals from the NCKU Ethics Committee Human Research, and all studies abided by the rules of the Internal Review Board of NCKU Hospital, IRB# A-ER-105-381. Informed consent both oral and written was obtained for all samples. All files are current and present.

### Histochemical staining

Whole mount alkaline phosphatase (ALP) staining was previously described^18^. Briefly, the skin was fixed with 4% paraformaldehyde (PFA) and then pre-incubated in ALP buffer overnight at 4°C. Then, it was incubated in ALP buffer with NBT/BCIP reagent at room temperature until color development and stopped with Tris-EDTA 20 mmol/L. After staining the whole mount tissue was cleared in benzyl alcohol benzyl benzoate (BABB).

Paraffin section alkaline phosphatase staining was previously described^18^. Briefly, 4% PFA-fixed paraffin-embedded sections were pre-incubated with ALP buffer for 1 hour at room temperature. Then, incubated in ALP buffer with NBT/BCIT reagent at room temperature until color development and stopped with Tris-EDTA 20 mmol/L. For trichrome staining, 4% PFA-fixed paraffin-embedded sections were incubated with Biebrich scarlet-acid fuchsin solution and Aniline blue solution (Masson’s Trichrome Stain kit, Sigma, St. Louis MO). Haematoxylin was used for nuclear counterstain. For H&E, the 4% PFA paraffin-embedded sections were fixed and then stained with haematoxylin and eosin (H&E) according to accepted protocol.

### Immunofluorescence staining

Tissues were collected from fetal and adult Lanyu pig, and either embedded in OCT for frozen sections or 4% PFA fixed overnight for paraffin sections. Pig tissues were cut into 7 μm sections using a microtome. For antigen retrieval, paraformaldehyde paraffin-embedded sections were incubated in sodium citrate in PBS pH 6 at 100°C for 20 minutes. The sections were rinsed and then blocked in 0.06% methanol peroxide and subsequently incubated with primary antibodies overnight at 4°C. Primary antibodies utilized; K5 at 1:500 (rabbit polyclonal, Abcam, Cambridge, UK), K10 at 1:500 (mouse monoclonal, Thermo, Waltham MA, USA), K17 at 1:500 (rabbit polyclonal, Abcam, Cambridge, UK), p63 at 1:200 (mouse monoclonal, Abcam, Cambridge, UK), PCNA at 1:200 (mouse monoclonal, Millipore, Burlington MA, USA), and Ctnnb1 at 1:200 (rabbit polyclonal, Sigma, St. Louis, MO, USA). The slides were washed multiple times and then incubated with secondary antibodies at room temperature for 1 hour (Alexa Fluor 488 IgG or Alexa Fluor 594 IgG Life Technologies, San Diego CA). Hoechst 33258 dye (Invitrogen, San Diego CA) was used for nuclear stain.

### Image acquisition

Gross images were documented with an Olympus OM-D hand-held camera. Microscopic images were recorded with either an Olympus BX51 Fluorescent Microscope or an Olympus FLUOVIEW FV1000 confocal microscope. Microscopic images of ALP+ cells were recorded with the 633λ (for reflectance of NBT/BCIP substrate) channel. Images in data figures are representative of at least 3 samples (n ≥ 3).

### Statistical analysis

All data figures are representative of at least three samples (n ≥ 3). Standard deviation calculation and 1-way ANOVA were performed on data sets to generate p-values for statistical significance. *P <0.05, **P <0.01, ***P <0.001.

## RESULTS

### Characterization of secondary hair follicle development in the Lanyu fetal pig

To characterize whether secondary hair follicle development in the Lanyu pig, we analyzed a dorsal skin developmental time course using alkaline phosphatase (ALP) staining (Fig. 1A–T).

**Figure 1.**
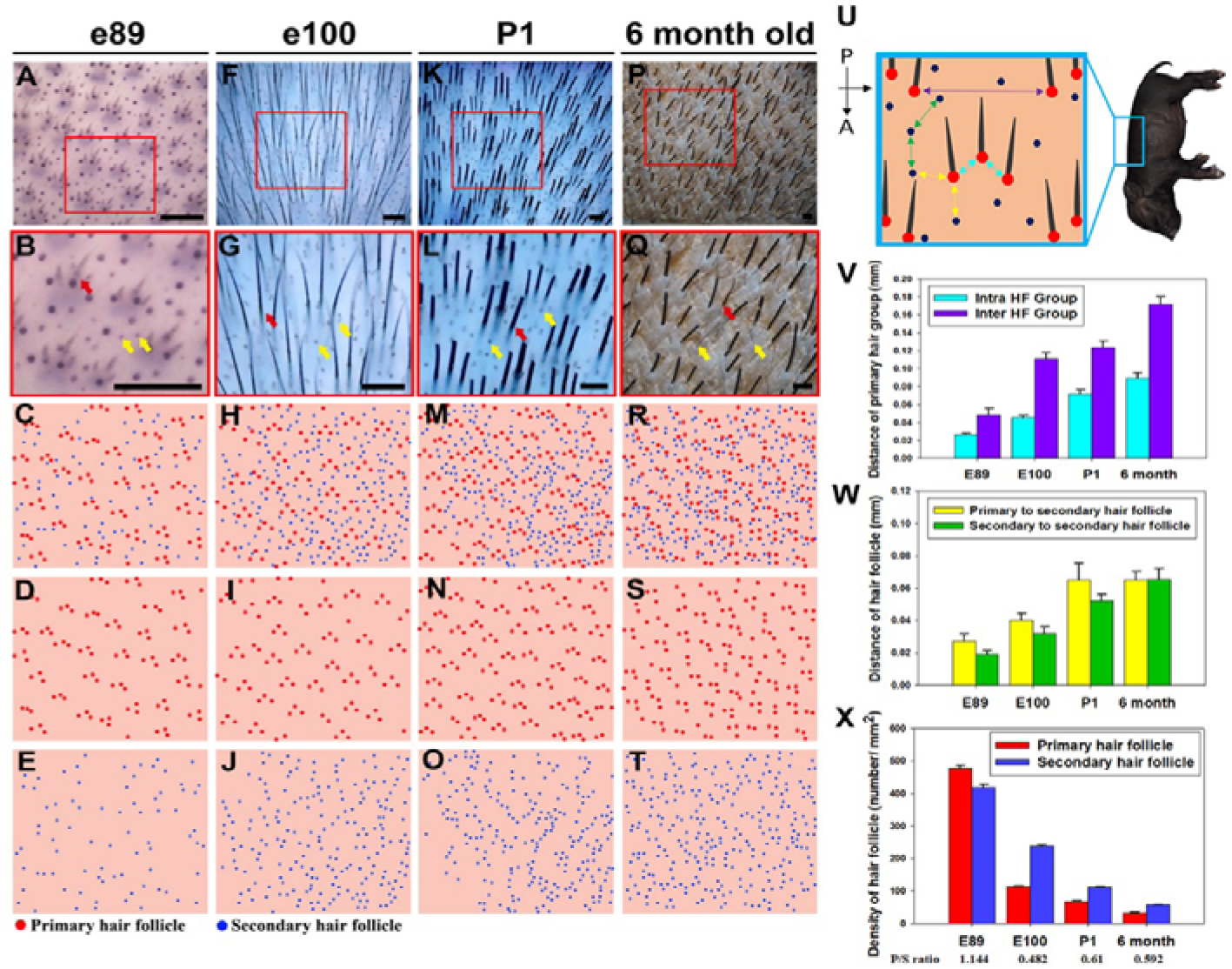
Spatiotemporal patterning of primary and secondary hair follicles in Lanyu pig dorsal skin. (A–T) Dorsal skin from Lanyu pigs was analyzed at (A-E) E89, (F-J) E100, (K-O) P1, and (P-T) 6 months. Alkaline phosphatase (ALP) staining identifies developing hair follicles at (A–B) E89, (F-G) E100, and P1 (K–O). Primary follicles (red) and secondary follicles (blue) were digitally annotated to illustrate spatial organization across developmental stages for (C-E) E89, (H-J) E100, (M-O) p1, and (R-T) 6 mo. (U) Schematic of distance measurements between follicle populations. (V) Inter- and intra-primary follicle distances. (W) Primary–secondary and secondary–secondary follicle distances and primary-to-secondary ratio. (X) Hair follicle density across developmental stages. (scale bar: 100 μm, red arrows: primary follicles; yellow arrows: secondary follicles, double-ended arrows: measured distances as labeled)

At E89, two distinct follicular populations were evident: (i) clusters of three large immature follicles arranged in a consistent spatial pattern, and (ii) smaller, ALP+, circular, punctate structures that were spatially independent of the large immature clusters (Fig. 1A–E). The clustered follicles exhibited a stable triplet organization, whereas the smaller structures appeared as isolated units.

By E100, both populations persisted but showed clear maturation differences. The triplet follicles increased in size and morphological complexity, while the smaller punctate, ALP+ structures began to exhibit emerging hair shafts and increased in number (Fig. 1F–J). Importantly, the spatial segregation between clustered and individual follicles was maintained.

At P1, the two populations remained clearly distinguishable, with mature large follicles retained in triplet clusters and smaller follicles distributed individually between clusters (Fig. 1K–O). This organization persisted into 6 months, where both follicle types exhibited mature morphology (Fig. 1P–T).

Due to the earlier emergence, larger size, and stable clustered organization, the triplet follicles were identified as primary hair follicles, whereas the later-developing, smaller, and individually distributed follicles were secondary hair follicles.

Spatially, primary follicles consistently formed chevron-like triplet patterns oriented along the anterior–posterior axis, while secondary follicles were randomly interspersed between these clusters throughout development.

Quantitative analysis revealed dynamic changes in follicle spacing and density (Fig. 1U–X). Both inter- and intra-primary follicle distances increased from E89 to 6 months (Fig. 1V), consistent with tissue expansion. Similarly, distances between primary–secondary and secondary–secondary follicles increased over time (Fig. 1W), while overall follicle density decreased (Fig. 1X). Notably, the primary-to-secondary follicle ratio decreased sharply between E89 and E100 and remained relatively stable thereafter, reflecting an increase of the secondary follicle population.

Taken together, these data show Lanyu pigs develop a distinct secondary hair follicle population emerging after primary follicle establishment and persisting into adulthood, forming a spatially organized pelage pattern.

### Whole-mount alkaline phosphatase staining defines stages of secondary hair follicle morphogenesis

To define the developmental progression of secondary hair follicles, we performed a whole-mount alkaline phosphatase (ALP) staining time course on Lanyu pig dorsal skin (Fig. 2A–I). ALP activity marks key follicular compartments, including placodes, epithelial sheaths, and dermal papillae, enabling visualization of follicle morphogenesis^18, 19^.

**Figure 2.**
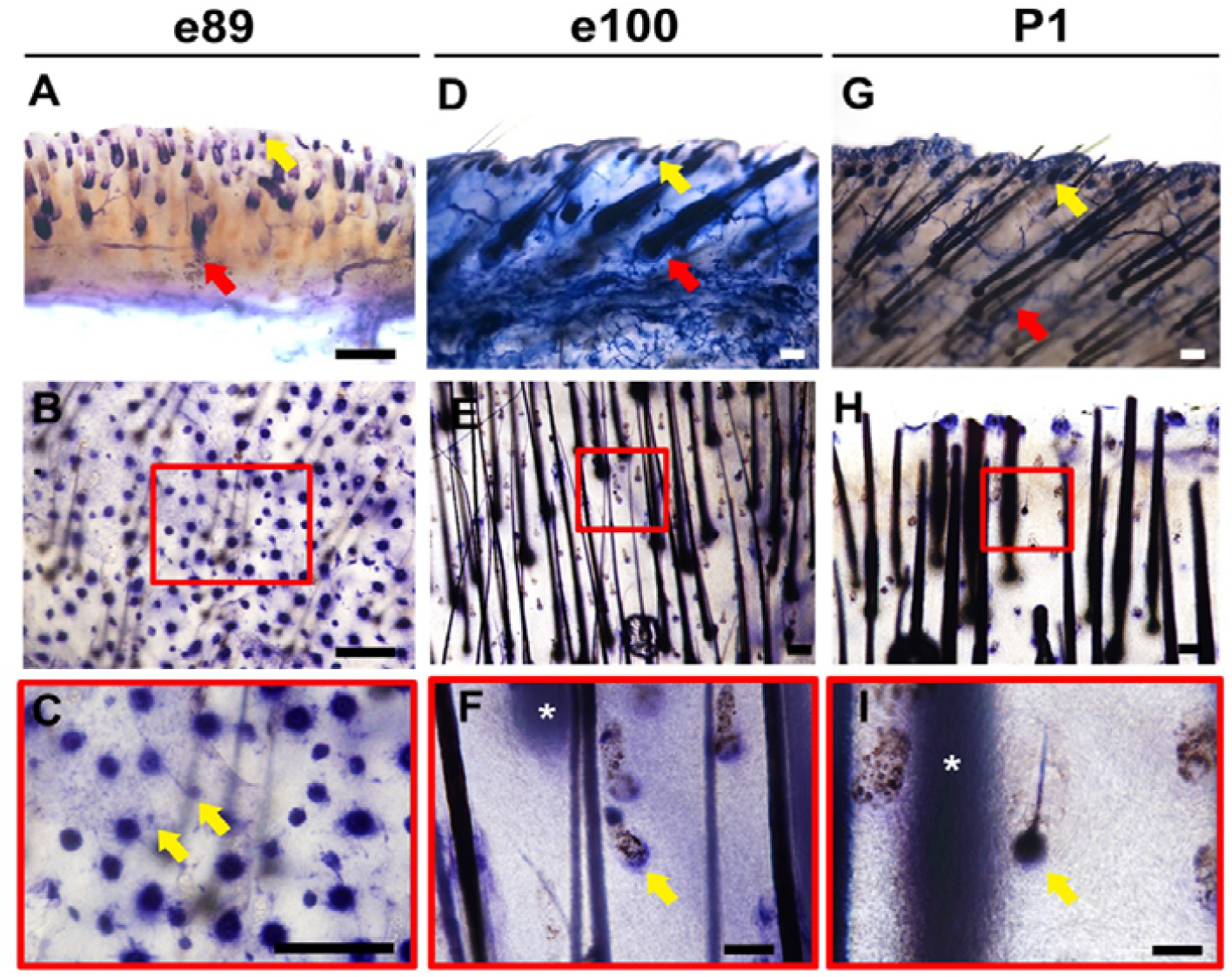
Whole-mount alkaline phosphatase staining reveals staged development of secondary hair follicles in Lanyu pig dorsal skin. (A–I) Whole-mount ALP staining of dorsal skin at E89, E100, and P1. (A–C) E89 skin showing primary follicles and secondary follicle placodes/hair germs. (D–F) E100 skin showing mature primary follicles and secondary hair pegs. (G–I) P1 skin showing mature primary and secondary follicles with visible hair shafts. For each stage, transverse sections (A, D, G) and top views (B–C, E–F, H–I) are shown. Primary follicles (red arrows) form clustered triplets, while secondary follicles (yellow arrows) are distributed between clusters. (scale bars: 100 μm, white asterisk (*): primary follicle out of focal plane)

At E89, secondary follicles were observed as placodes and early hair germs, appearing as small ALP+ structures spatially interspersed between primary follicle triplets (Fig. 2A–C). These placodes and hair germs were consistently excluded from within the primary follicle clusters, establishing an early spatial segregation between follicle types.

By E100, secondary follicles progressed to the hair peg stage, characterized by elongated epithelial downgrowth and increased ALP signal intensity (Fig. 2D–F). At this stage, primary follicles were morphologically mature, while secondary follicles were immature, remained smaller and continued to localize between primary hair triplets.

At P1, secondary follicles developed full maturation, exhibiting well-defined follicular architecture and visible hair shafts (Fig. 2G–I). Despite maturation, secondary follicles remained markedly smaller than primary follicles and maintained their inter-cluster spatial distribution.

We observed four sequential stages of secondary hair follicle development in the Lanyu pig: placode → hair germ → hair peg → mature follicle. Importantly, this progression occurs after the establishment of primary follicle triplets and is spatially restricted to the interstitial regions between them.

Together, these data show secondary hair follicles developed through a conserved morphogenetic sequence, with delayed timing and distinct spatial positioning relative to primary follicles, ultimately forming an integral component of the neonatal pelage.

### Secondary hair follicle morphogenesis exhibits conserved protein expression patterns

To characterize the molecular and cellular features of secondary hair follicle development, paraffin sections from defined developmental stages were immunostained for basal or suprabasal keratinocyte markers (K5, K10), hair keratin (K17), morphogen signaling (β-catenin), and hair follicle development marker (ALP) (Fig. 3; S1 Fig).

**Figure 3.**
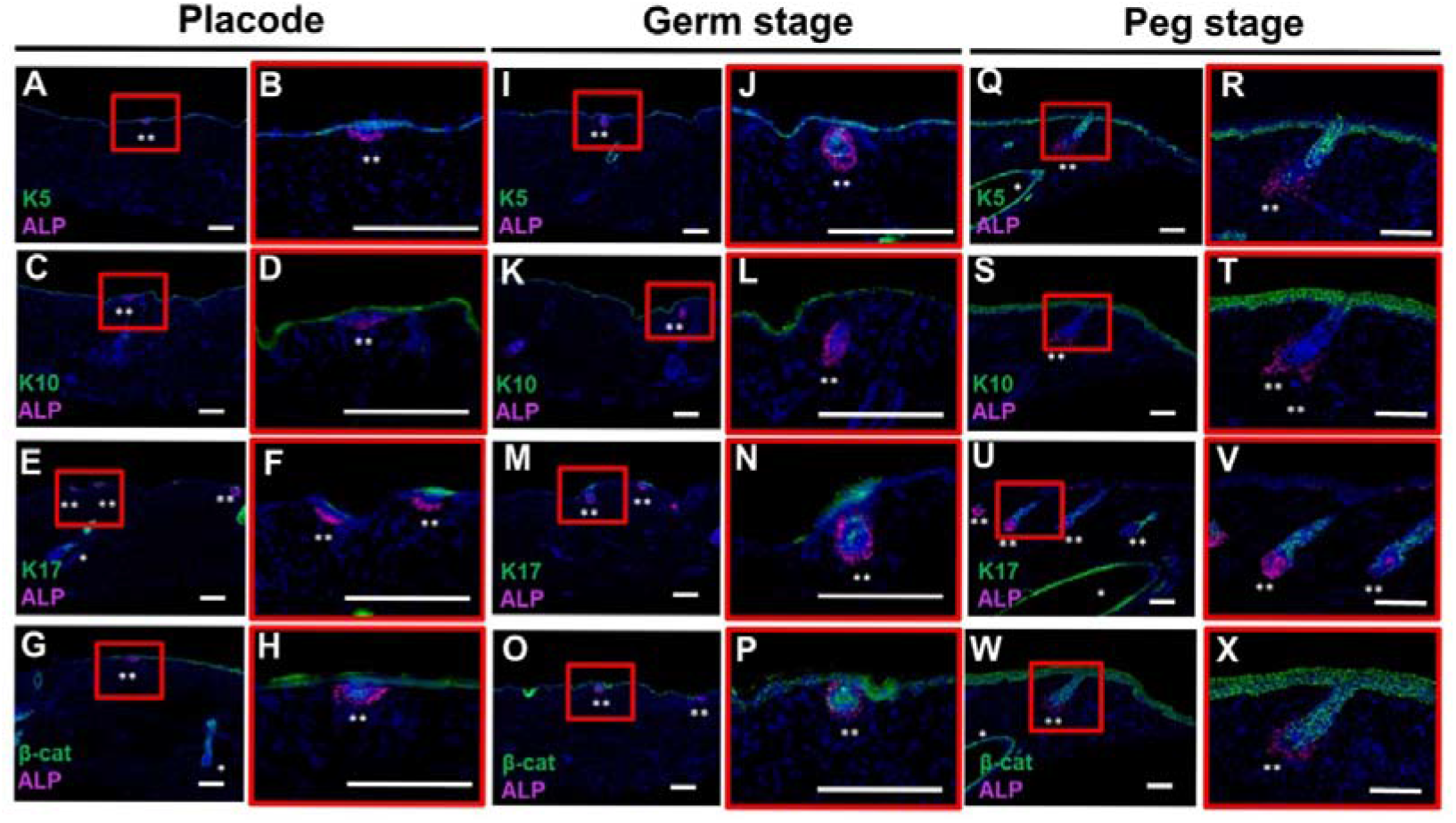
Spatiotemporal protein expression patterns during secondary hair follicle development in Lanyu pig dorsal skin. (A–X) Paraffin sections of secondary hair follicles at placode, hair germ, and hair peg stages immunostained for K5, K10, K17, and β-catenin, followed by alkaline phosphatase (ALP) staining to identify dermal papilla cells. (A–H) Placode stage: K5 (A–B), K10 (C–D), K17 (E–F), β-catenin (G–H). (I–P) Hair germ stage: K5 (I–J), K10 (K–L), K17 (M–N), β-catenin (O–P). (Q–X) Hair peg stage: K5 (Q–R), K10 (S–T), K17 (U–V), β-catenin (W–X). K5 marks basal keratinocytes, K10 marks suprabasal differentiation, and K17 highlights follicular epithelium. (scale bars: 100 μm, blue: Hoechst, white asterisks (*): primary follicles, white asterisks (**): secondary follicles)

At the placode stage, the epidermis consisted of a single K5+ basal layer with localized epithelial thickening overlying ALP+ mesenchymal condensates (Fig. 3A– B). K10□ suprabasal cells were present but remained spatially separated from the mesenchyme (Fig. 3C–D). In contrast, K17+ epithelial cells formed a distinct cluster directly over ALP+ mesenchymal cells, marking early follicular specification (Fig. 3E–F). These epithelial–mesenchymal interfaces were enriched for nuclear β-catenin+ cells, consistent with active Wnt signaling during placode formation (Fig. 3G–H). Compared to interfollicular epidermis, placodes exhibited increased epithelial stratification and proliferation via p63+ and PCNA+ cells respectively (S1 Fig).

During the hair germ stage, epithelial downgrowth was observed, with expanded K5+ basal cells forming an invaginating structure that maintained direct contact with an enlarging ALP+ dermal condensate (Fig. 3I–J). K10+ cells remained restricted to suprabasal layers and maintained separation from the mesenchyme (Fig. 3K–L). K17+ cells increased in number and remained concentrated over the ALP+ mesenchymal cells, while nuclear β-catenin activity persisted in epithelial cells adjacent to these ALP+ cells (Fig. 3M–P). These changes were accompanied by further epithelial stratification (p63+) and increased proliferative (PCNA+) activity (S1 Fig).

At the hair peg stage, a well-defined epithelial column extended into the dermis, composed of K5+ basal cells forming a polarized downgrowth in continuous contact with a now-expanded and deeper-positioned ALP+ dermal papilla (Fig. 3Q–R). K10+ suprabasal cells increased in number and formed multiple layers above the basal compartment and further remote from the ALP+ dermal cells (Fig. 3S–T). K17 expression became strongly enriched along the elongating follicular epithelium, particularly in regions adjacent to the dermal papilla (Fig. 3U–V). Notably, β-catenin localization shifted predominantly to cell–cell junctions with reduced nuclear signal, suggesting a transition in Wnt signaling dynamics during follicle maturation (Fig. 3W–X). At this stage, size differences between primary and secondary follicles were clearly apparent (Fig 3W and S1 Fig).

Collectively, these data demonstrate Lanyu pig secondary hair follicles undergo a canonical morphogenetic progression characterized by epithelial stratification, sustained epithelial–mesenchymal interaction, and dynamic β-catenin signaling.

### Histological reconstruction defines a developmental timeline of secondary hair follicle morphogenesis

To integrate the molecular and whole-mount findings into a unified developmental framework, we performed histological analysis utilizing hematoxylin and eosin (H&E) staining of Lanyu pig fetal dorsal skin across key timepoints (Fig. 4A–F).

**Figure 4.**
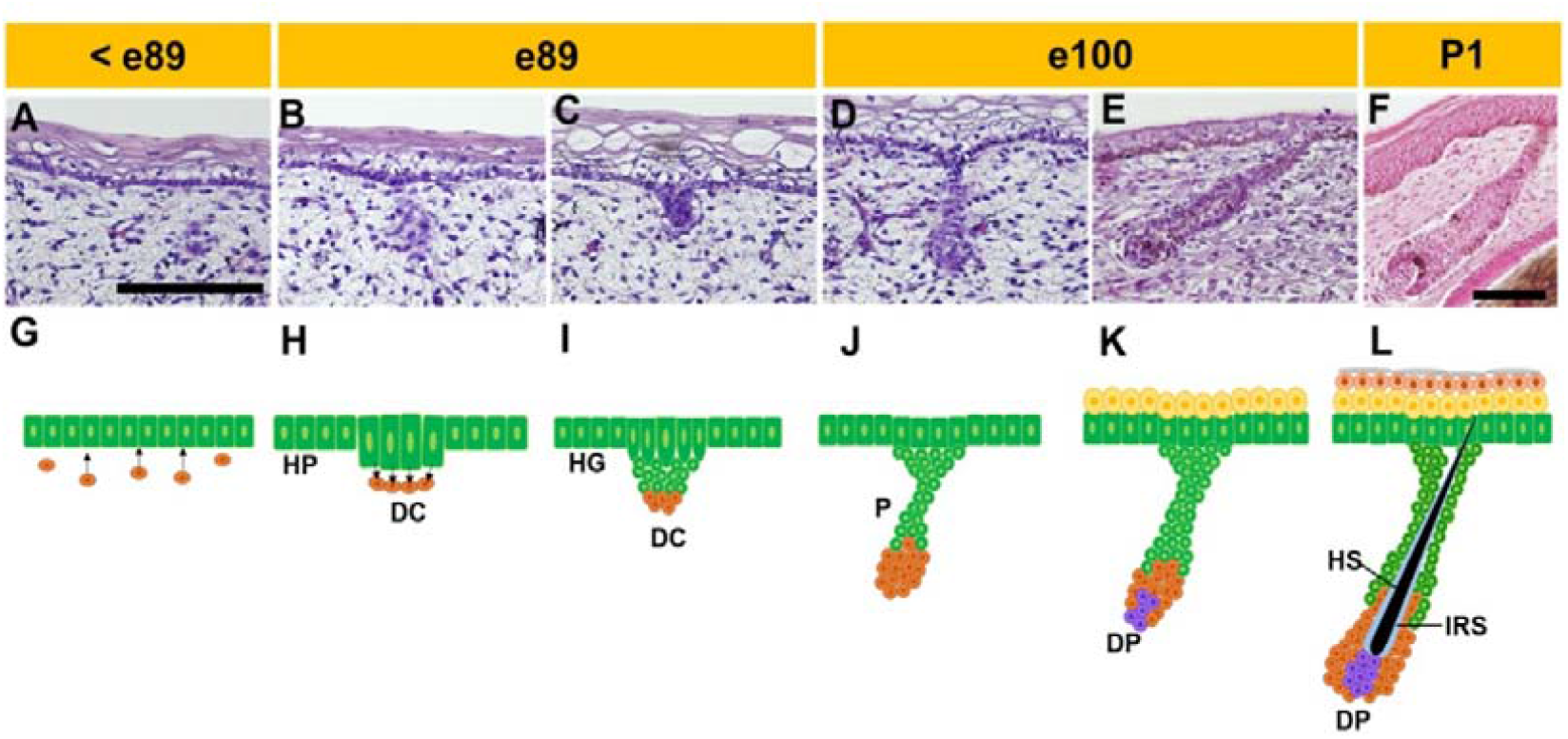
Histological reconstruction of secondary hair follicle morphogenesis in Lanyu pig dorsal skin. (A–F) Hematoxylin and eosin (H&E) staining of dorsal skin paraffin sections across developmental stages. (A) Pre-placode epidermis showing a single epithelial layer covered by periderm. (B–C) Placode stage with epithelial thickening and underlying dermal condensate. (C–D) Hair germ stage showing epithelial invagination and expanded dermal condensate. (D–E) Hair peg stage with elongated epithelial downgrowth partially surrounding the dermal papilla. (F) Mature secondary follicle at P1 displaying infundibulum, hair shaft, matrix, and dermal papilla. (G–L) Schematic model summarizing secondary hair follicle developmental stages. (DC: dermal condensate, DP: dermal papilla, HP: hair placode, HG: hair germ, P: hair peg, HS: hair shaft, IRS: inner root sheath, scale bars: 100 μm)

Prior to E89, the epidermis consisted of a single epithelial layer covered by a periderm, with no evidence of follicular initiation or underlying mesenchymal condensation (Fig. 4A).

At approximately E89, localized epithelial thickenings emerged, forming secondary hair placodes characterized by clusters of columnar basal cells overlying nascent dermal condensates (Fig. 4B–C).

From E89 to E100, progressive epithelial stratification was accompanied by epithelial invagination into the dermis, while dermal cells condensed into a defined structure beneath the epithelium, consistent with the hair germ stage (Fig. 4C–D).

By E100, continued epithelial downgrowth resulted in the formation of hair pegs, in which elongated epithelial cells partially enveloped the dermal condensate, establishing a more mature epithelial–mesenchymal unit (Fig. 4D–E).

At P1, secondary follicles exhibited fully developed architecture, including the infundibulum, hair shaft, matrix, and dermal papilla (Fig. 4F; S1 Fig.). At this stage, the interfollicular epidermis displayed regularly spaced epithelial indentations, but fully developed rete ridges were absent, consistent with previous reports that rete ridge formation occurs postnatally in Lanyu pig skin.

Utilizing these observations, we constructed a developmental model summarizing secondary hair follicle morphogenesis (Fig. 4G–L), encompassing the sequential stages of placode formation, hair germ development, and hair peg maturation, culminating in a structurally complete follicle at birth.

Collectively, these data help establish a temporal and morphological framework for secondary hair follicle development in the Lanyu pig, consistent with canonical folliculogenesis, while defining species-specific developmental timing and epidermal context.

### Lanyu pig secondary hair follicle exhibits structural and molecular similarity to human vellus hair

To assess translational relevance, we compared mature Lanyu pig secondary follicles with human vellus hair follicles (Fig. 5). Paraffin sections from Lanyu pig and human skin were stained with H&E and immunostained for basal or suprabasal keratinocyte markers (K5, K10), and hair follicle stem cell marker cytokeratin 15 (K15) (Fig. 5).

**Figure 5.**
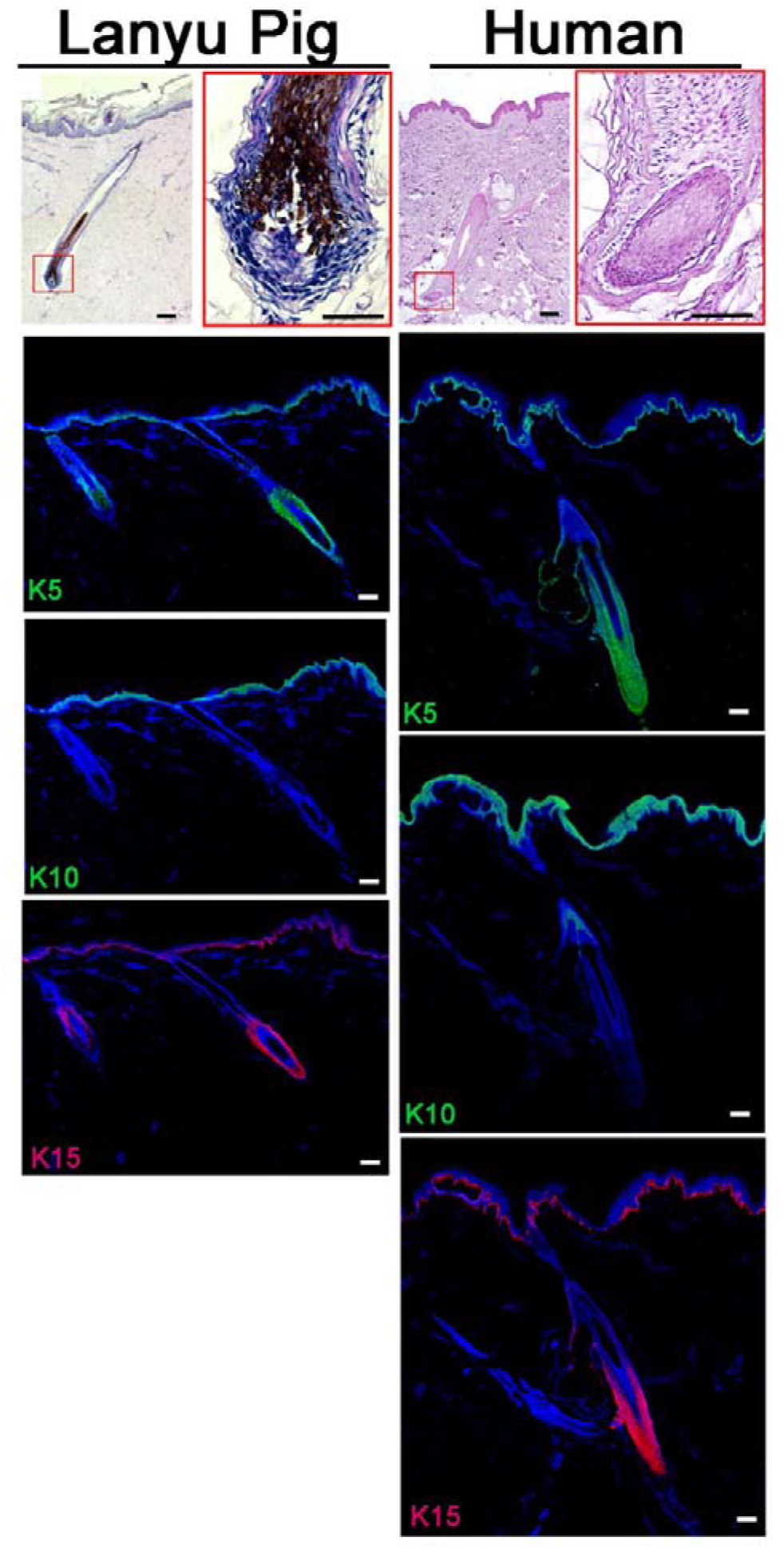
Structural and molecular similarity between Lanyu pig secondary hair and human vellus hair. Comparative analysis of Lanyu pig secondary hair follicles and human vellus hair follicles. Hematoxylin and eosin (H&E) staining demonstrates similar follicle size, morphology, and dermal papilla organization between species. Immunostaining shows conserved expression patterns of cytokeratin 5 (K5) in basal keratinocytes and cytokeratin 10 (K10) in suprabasal layers in both follicular and interfollicular epidermis. Cytokeratin 15 (K15) marks the hair follicle stem cell niche (bulge region) in both Lanyu pig and human follicles. (scale bars: 50 μm, blue: Hoescht)

Histological analysis revealed comparable dermal papilla and matrix organization (Fig 5 top panels), and conserved spatial expression patterns of K5 and K10 within basal and suprabasal compartments, respectively (Fig. 5 middle panels). Additionally, the stem cell marker K15 localized to the bulge region in both species, indicating conservation of follicular stem cell location and hair follicle architecture (Fig. 5 bottom panels).

Collectively, these data suggest the mature Lanyu pig secondary hair follicle structure closely resembles human vellus hair follicles architecture.

## DISCUSSION

In this study, we define the development, patterning, and molecular characteristics of secondary hair follicles in the Lanyu pig and establish this system as a translational model for human vellus hair biology. Our findings demonstrate that Lanyu pigs not only develop but retain secondary hair follicles into adulthood, in contrast to commonly used porcine models in which these structures regress.

We show that secondary follicles arise following primary follicle formation, occupy distinct interfollicular positions, and exhibit reduced size and density relative to primary follicles (Fig. 1-2). These follicles undergo a canonical morphogenetic sequence—placode, hair germ, and hair peg—accompanied by conserved epithelial–mesenchymal interactions and dynamic β-catenin signaling (Fig. 3-4)^4^. Importantly, the structural organization and protein expression patterns of Lanyu pig secondary follicles closely resemble those of human vellus hair follicles, supporting their translational relevance (Fig. 5)^2, 20-22^.

A key implication of this work is the identification of a previously underutilized porcine model that preserves secondary hair follicle biology. In Landrace pigs, Meyer and Gorgen showed secondary follicles initiate but regress during late fetal or early postnatal development, limiting their applicability for studies of hair follicle–mediated regeneration^3^. In contrast, the persistence of secondary follicles in the Lanyu pig provides a unique opportunity to investigate the role of vellus-like follicles in skin homeostasis and repair (Fig. 1-5).

These findings are particularly relevant in the context of wound healing. In humans, LFT injuries result in scarring without regeneration of vellus hair follicles, whereas rodent models exhibit partial regeneration limited to secondary follicles ^15, 16, 23-27^. Notably, only rare species such as the African spiny mouse demonstrate complete regeneration of both primary and secondary follicles, but these animals possess loose skin and therefore differ fundamentally from human physiology^17, 28^. The Lanyu pig, as a tight-skinned mammal with preserved secondary follicles, fills a critical gap between existing models and human biology.

The location of interfollicular skin stem cells have been reported in the basal layer ^9^, the bottom of the rete ridge basal layer ^29-31^, and the top of the rete ridge basal layer ^31-33^. Intuitively, the position of these skin stem cells at the bottom of the rete ridge offers protection of this population from surface trauma but requires further study. Humans fail to regenerate rete ridges after LFT wounding^34^. Lanyu pig skin architecture shows the presence of rete ridges (Fig. 4-5). In addition, ALP staining in human and Lanyu pig skin showed positive cells at the bottom of the rete ridge basal layer suggesting a stem cell niche ^18^. Both species ALP+ cells were K5+, K10-, and expressed less PCNA than inter-rete ridge basal cells. Collectively, this knowledge further supports Lanyu pig skin as a translational model for human LFT wound research.

Hair follicle stem cells play a central role in regenerative responses by contributing to re-epithelialization and tissue repair^8, 35-42^. The absence of secondary follicles in scar tissue reduces the available HFSC pool, limiting regenerative capacity and predisposing tissue to functional decline following repeated injury. Our findings suggest that restoring or preserving vellus-like follicles (Fig. 5) could represent a key strategy for improving regenerative outcomes in human skin.

In addition to developmental characterization, we provide evidence that the secondary hair follicle trait in the Lanyu pig follows an autosomal dominant inheritance pattern, as demonstrated by its retention in hybrid offspring (Sup Fig. 2-3). This observation suggests that secondary follicle formation represents an ancestral trait that has been selectively lost in modern commercial breeds. Most domestic pigs, not originating from Timor or Papua New Guinea, are derived from wild boars and grouped into four regions; Western, Eastern, Indian, and Indonesian ^43^. Wild boars develop and retain secondary hair follicles ^44, 45^. Lanyu pigs (*Sus scrofa taivanus*) are domesticated from the Taiwanese wild boar (*Sus scrofa taivanus*) indigenous to Taiwan ^46, 47^. The identification of genetic determinants underlying this phenotype represents an important avenue for future investigation and may enable targeted breeding or genetic approaches to restore secondary follicle development in other models.

Finally, our study provides a foundation for future functional work. Integrating wound healing models with transcriptomic and single-cell analyses in the Lanyu pig will enable dissection of the molecular pathways governing ectodermal organ regeneration. In particular, defining the behavior and niche dynamics of hair follicle stem cells during injury and repair will be critical for translating these findings into therapeutic strategies.

In summary, we establish the Lanyu pig as a novel and clinically relevant model for secondary (vellus-like) hair follicle biology, bridging a critical gap between traditional animal models and human skin. This system provides a platform for investigating the mechanisms limiting ectodermal organ regeneration and for developing strategies to enhance regenerative healing in human patients.

## CONCLUSION

In summary, the Lanyu pig develops and retains pelage secondary hair follicles into adulthood, distinguishing it from commonly used porcine models in which these structures regress. These follicles arise through a canonical morphogenetic sequence—placode, hair germ, and hair peg—driven by coordinated epithelial–mesenchymal interactions and conserved molecular signaling pathways.

Importantly, Lanyu pig secondary follicles closely resemble human vellus hair follicles in both structure and protein expression, supporting their relevance as a translational model. The observation that this trait follows an autosomal dominant inheritance pattern further suggests that secondary follicle development represents an ancestral characteristic that has been lost in modern commercial breeds.

Collectively, these findings establish the Lanyu pig as a clinically relevant, tight-skinned model that preserves vellus-like hair follicles, providing a valuable platform for investigating mechanisms of skin regeneration and for developing strategies to restore ectodermal organ function following injury.

## ACKNOWLEDGEMENT

The authors wish to thank Professor Cheng-Ming Chuong MD PhD in the Department of Pathology, University of Southern California, Los Angeles, California, USA for his discussion and insight.

## Author Contributions

MWH and CHL conceived the idea. CHL, YYH, and MWH carried out experiments. TYK and LHH provided study materials. CHL, CCW, MJT, SJS, and MWH analyzed the data. CHL and MWH wrote the manuscript.

## FIGURES

**S1 Fig.**
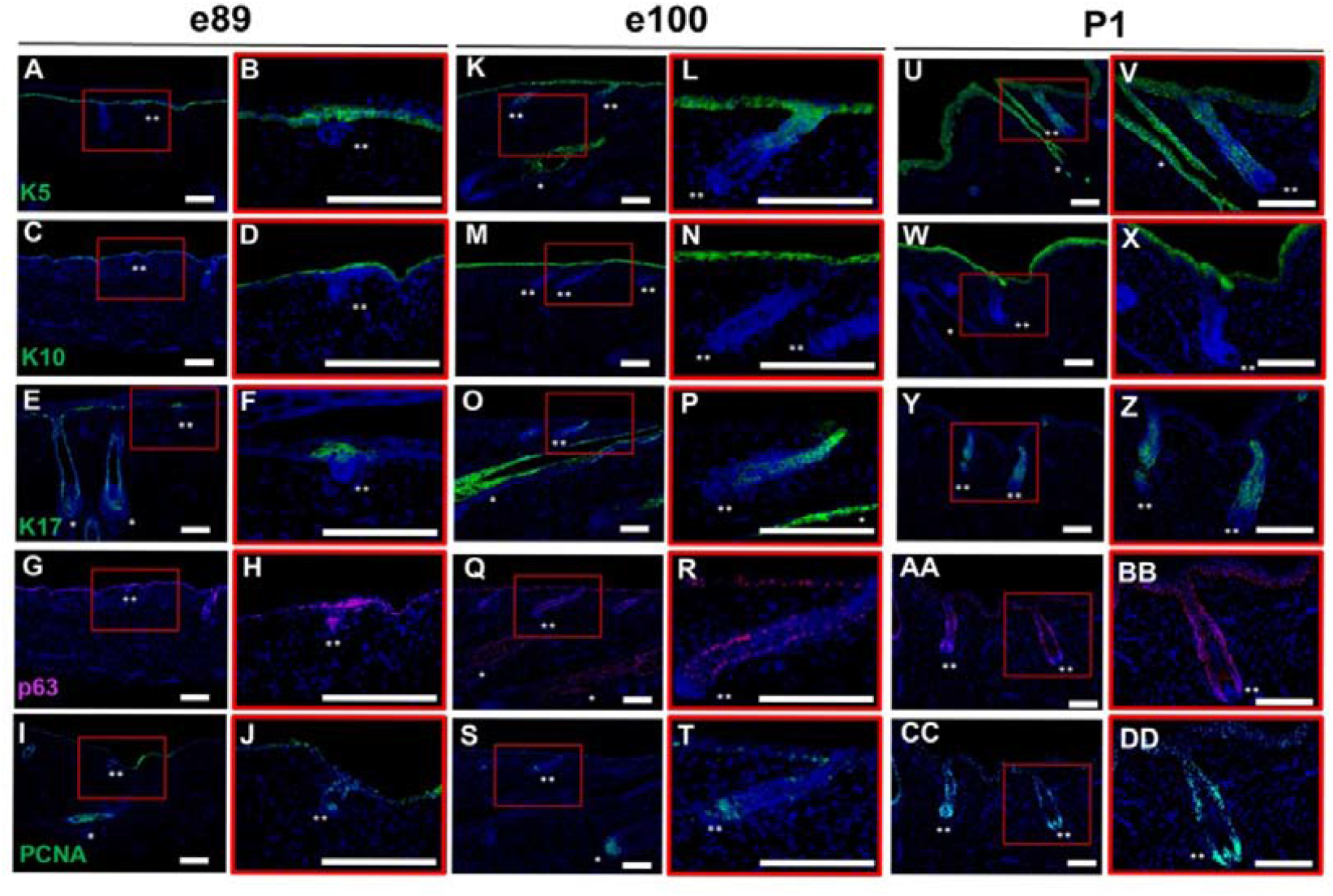
Spatiotemporal protein expression patterns during Lanyu pig hair follicle development. Paraffin sections of Lanyu pig dorsal skin were immunostained to assess protein expression patterns of K5, K10, K17, TRP63, and PCNA during hair follicle development across stages. (A–J) Secondary hair placode and hair germ stages at E89. (K–T) Secondary hair peg and early follicle stages at E100. (U–DD) Primary and secondary hair follicles at P1. (scale bars: 100 μm, blue: Hoecsht, white asterisks (*): primary follicles, double white asterisks (**): secondary follicles)

**S2 Fig.**
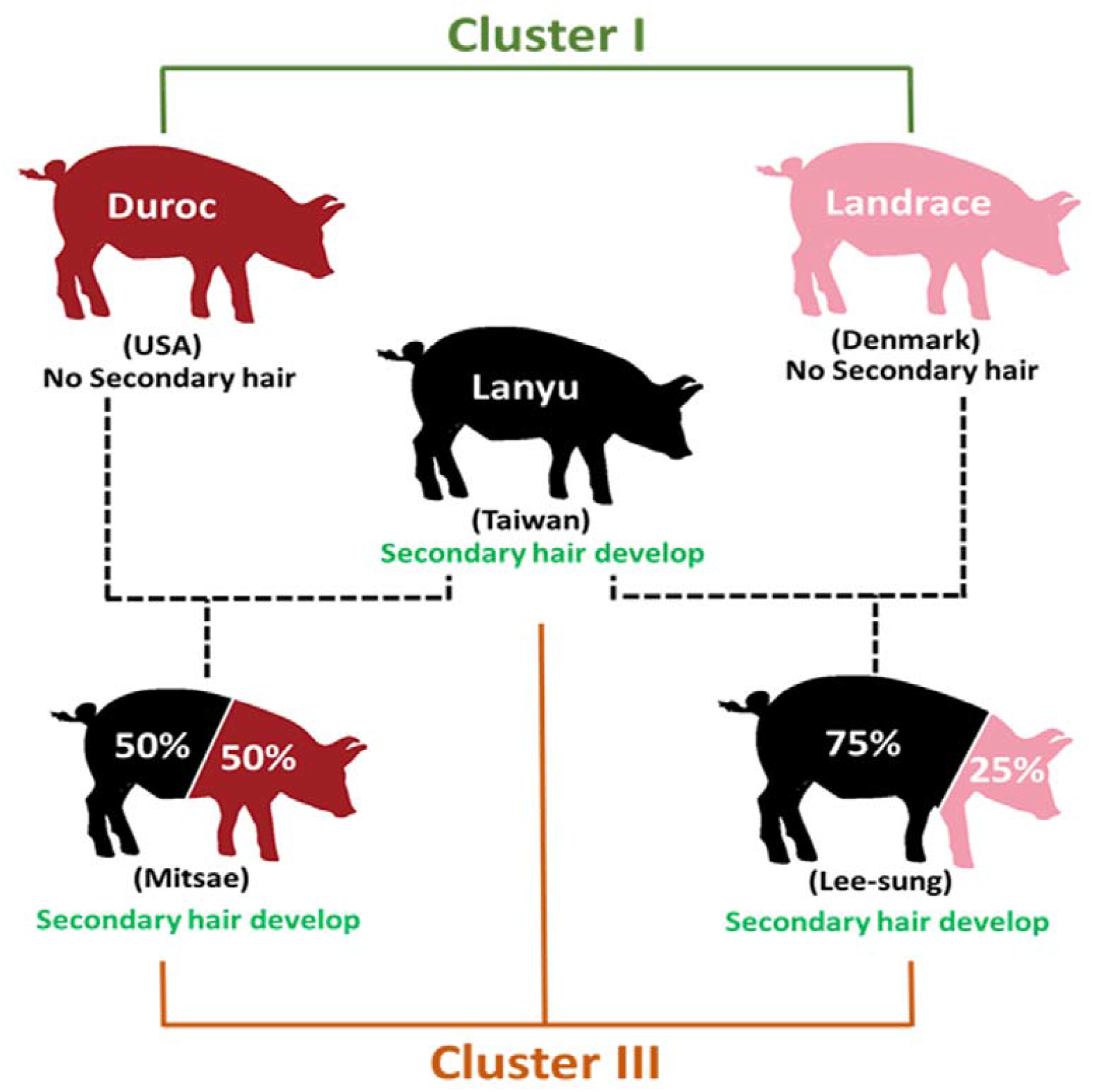
Secondary hair follicle trait and genetic relationships among pig breeds. Comparison of Lanyu pig pelage secondary hair follicle retention compared to Red Duroc and Landrace pig breeds. Related genetic background information is overlayed from Li et al., 2014. Red Duroc and Landrace pigs do not retain secondary hair follicles into adulthood, whereas Lanyu pigs maintain these structures. Lanyu-derived outcrosses, including Mitsai and Lee-Sung pigs, also retain secondary hair follicles, supporting dominant inheritance of this trait. (red: Red Duroc, pink: Landrace, black: Lanyu)

**S3 Fig.**
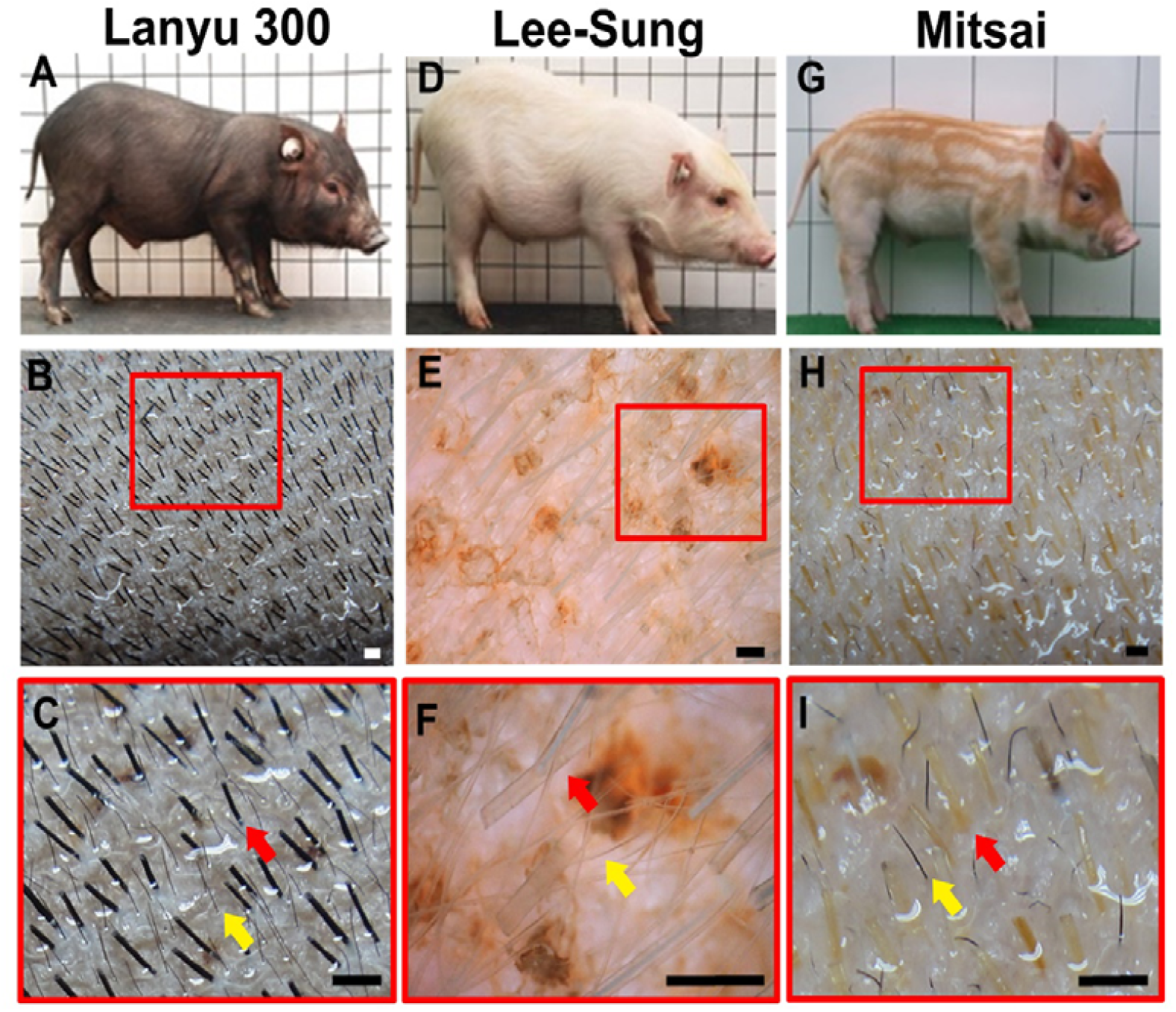
Inheritance of secondary hair follicles in Lanyu pig outcrosses. Gross dorsal skin images illustrating hair follicle patterning in Lanyu pigs and outcrossed strains. (A–C) Lanyu pig. (D–F) Lee-Sung pig (Lanyu × Landrace). (G–I) Mitsai pig (Lanyu × Red Duroc). Both outcrossed strains retain secondary hair follicles, consistent with an autosomal dominant inheritance pattern. (scale bars: 100 μm, red arrows: primary follicles, yellow arrows: secondary follicles, images in (A), (D), and (G) adapted from the Livestock Research Institute, Council of Agriculture, Executive Yuan, Taiwan)

