## Supplementary figures and images for "Spatiotemporal secondary hair follicle development in the Lanyu pig (Sus scrofa taivanus): a novel pelage hair follicle model"

### Supplemental Figure 1

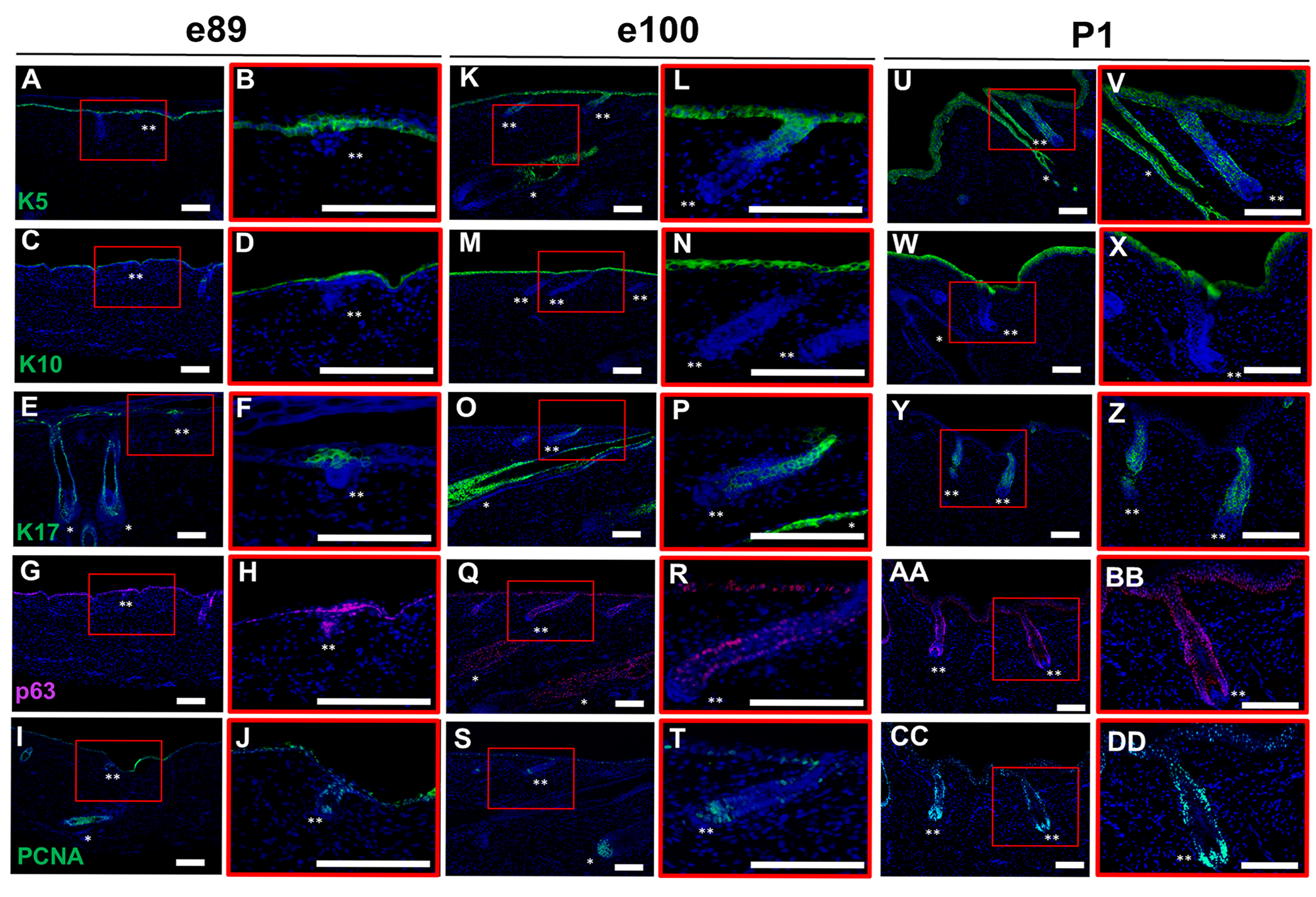

### Supplemental Figure 2

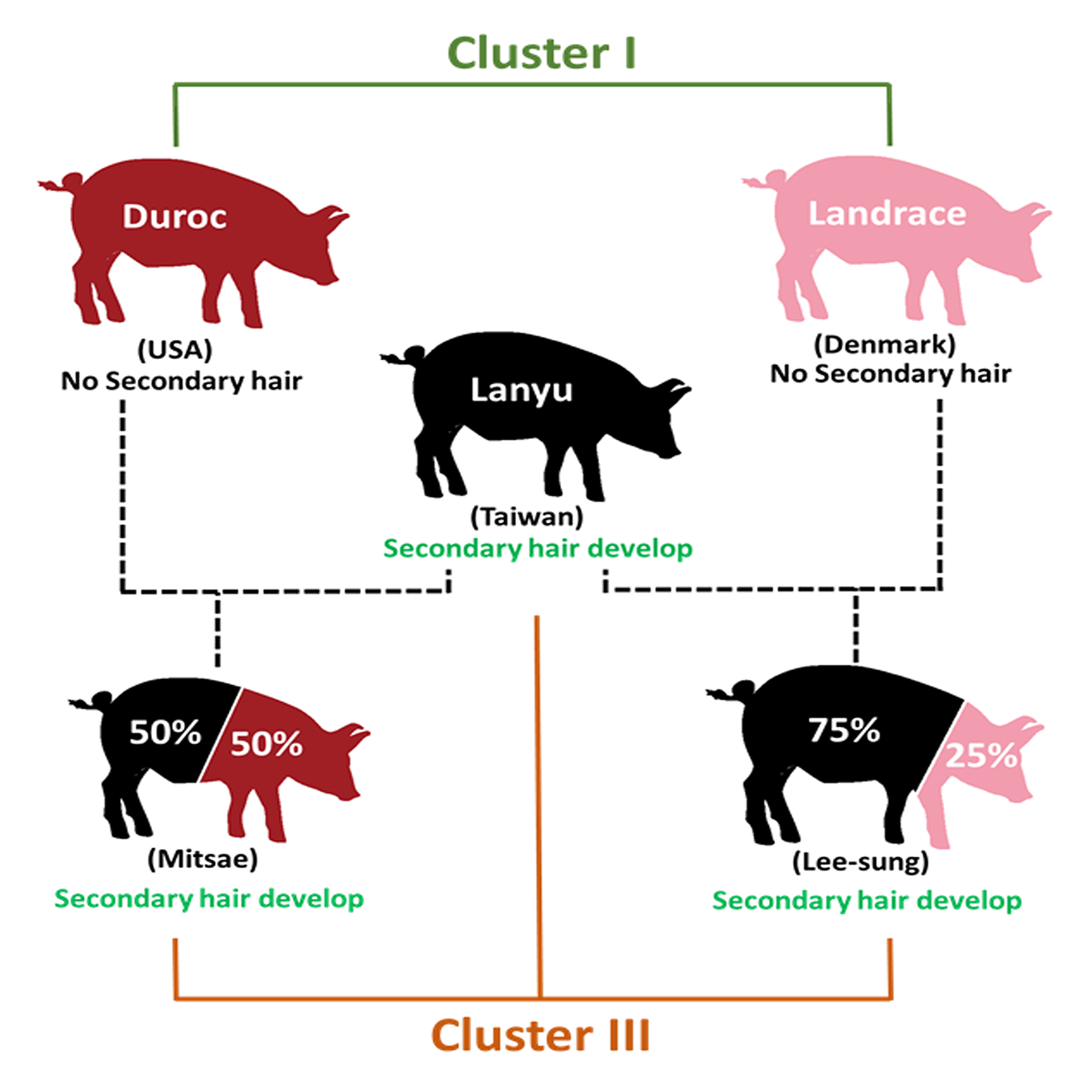

### Supplemental Figure 3

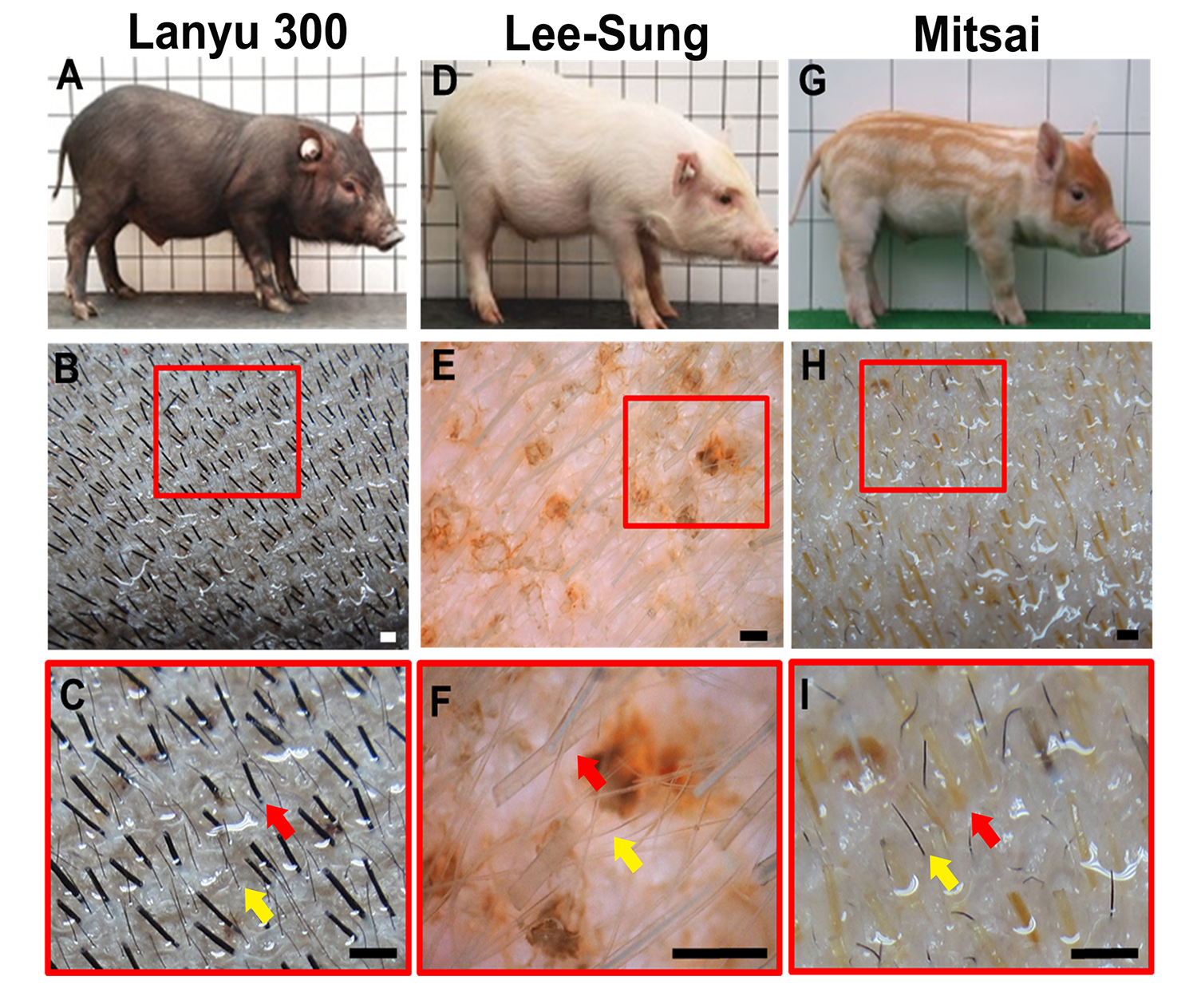
